# Epithelial mechanics in a three-dimensional cell vertex model with local junctional constraints

**DOI:** 10.64898/2026.05.15.725329

**Authors:** Kodai Terada, Yohei Kondo

## Abstract

Epithelial monolayers must accommodate deformation while maintaining continuous tissue coverage, but how their material properties emerge from three-dimensional cell mechanics remains incompletely understood. Among several three-dimensional cell vertex formulations, we build on a minimal model that captures experimentally observed features of epithelial monolayers, including shape bistability. The geometric constraint introduced in the model, however, is limited to regular cells undergoing uniform deformation. Here, we extend this model by replacing the original geometric constraint with a local energy based on the geometry of cell-cell junctions, allowing general vertex motions while retaining analytical tractability for regular cell packings and enabling simulations of disordered tissues. Using this framework, we derive analytical expressions for the in-plane elastic moduli and Poisson’s ratio and characterize the shear modulus across the tissue thickness. We show that the model reproduces a near-zero Poisson’s ratio observed in cultured epithelia. Numerical simulations show that this response persists beyond uniform deformation and remains robust to structural disorder over the timescales examined. Structural disorder also broadens the spectrum of internal relaxation modes and thus introduces slower modes to the frequency-dependent mechanical response. Together, these results establish a tractable three-dimensional framework for understanding how epithelial tissue mechanics emerges from cell-level geometry and interactions under realistic deformations and structural disorder.

## INTRODUCTION

Material properties of biological tissues play essential roles in morphogenesis and physiological function [1, 2]. Epithelial tissues, such as the Drosophila wing disc and cultured epithelial monolayers, provide accessible model systems for quantifying tissue viscoelasticity [3–5]. These studies have revealed material properties characteristic of living tissues. One notable example is their small in-plane Poisson’s ratio [6, 7], indicating little transverse contraction under tensile stretching. In parallel, substantial effort has been devoted to characterizing the mechanical properties of individual cells using a variety of experimental techniques [8, 9]. However, how tissue-scale material properties emerge from cellular-scale mechanics remains incompletely understood, representing an important question in biophysics.

Cell vertex models (CVMs) are theoretical tools for studying the relationship between cell mechanics and tissue mechanics [10–13]. CVMs represent each cell as a polygon or polyhedron, with vertex positions determined by an energy function that reflects cellular mechanical interactions. Their continuum-level elastic and viscoelastic properties have also been investigated [14, 15]. While two-dimensional CVMs have been studied most extensively, three-dimensional formulations can capture mechanical effects associated with apico-basal architecture and volumetric cell geometry that are absent from planar models [16–20]. At the same time, three-dimensional models admit a broader range of constitutive choices, and their predictions can depend sensitively on the modeling assumptions [21]. This makes experimentally motivated constraints particularly important when constructing three-dimensional CVMs.

Among these models, the framework introduced by Hannezo and colleagues provides a minimal description of epithelial mechanics based on three-dimensional cell geometry [17]. As we show below, this framework gives rise to a near-zero in-plane Poisson’s ratio over a broad region of parameter space, consistent with mechanical measurements of epithelial monolayers [6, 7]. This contrasts with two-dimensional cell vertex models, in which a zero Poisson’s ratio occurs only for a restricted set of parameters [22, 23]. However, the geometric confinement in the original three-dimensional formulation is defined for regular prismatic cells and does not extend naturally to general non-affine vertex motions. This limitation is particularly relevant when considering disordered tissues, where deformation can involve substantial non-affine vertex motions [21].

Here, we therefore extend the original model to describe non-affine relaxation. To this end, we replace its global geometric confinement with a local energy defined by the configuration of edges meeting at each cell corner. This extension allows us to analytically derive tissue-scale elastic moduli and Poisson’s ratio in the infinitesimal-strain regime, while also enabling numerical computation of the complex moduli of disordered tissues.

Using this framework, we first analyze the in-plane elasticity of regular prismatic cells and show that the model exhibits a near-zero Poisson’s ratio over a broad parameter range. Although the analytical result is derived in the infinitesimal-strain limit, it remains in good agreement with numerical calculations under finite strains, including biologically relevant deformations. We then examine mechanical responses that are specific to the three-dimensional formulation. In particular, regular prismatic cells can lose stability against out-of-plane simple shear, leading to stable configurations at finite shear. Finally, we numerically investigate the frequency-dependent viscoelastic response and the contributions of non-affine relaxation modes. Structural disorder broadens the relaxation spectrum and enhances the contribution of slow modes to the low-frequency response.

## MODEL

We build on the three-dimensional vertex model introduced previously and augment it with a local vertex-based regularization that suppresses degenerate vertex configurations. Cells are modeled in three dimensions (Figure 1a) and densely packed without gaps, representing a confluent monolayer tissue (Figure 1c). Cells are treated as incompressible in three dimensions, an assumption supported experimentally in systems such as MDCK monolayers [6].

**FIG. 1.**
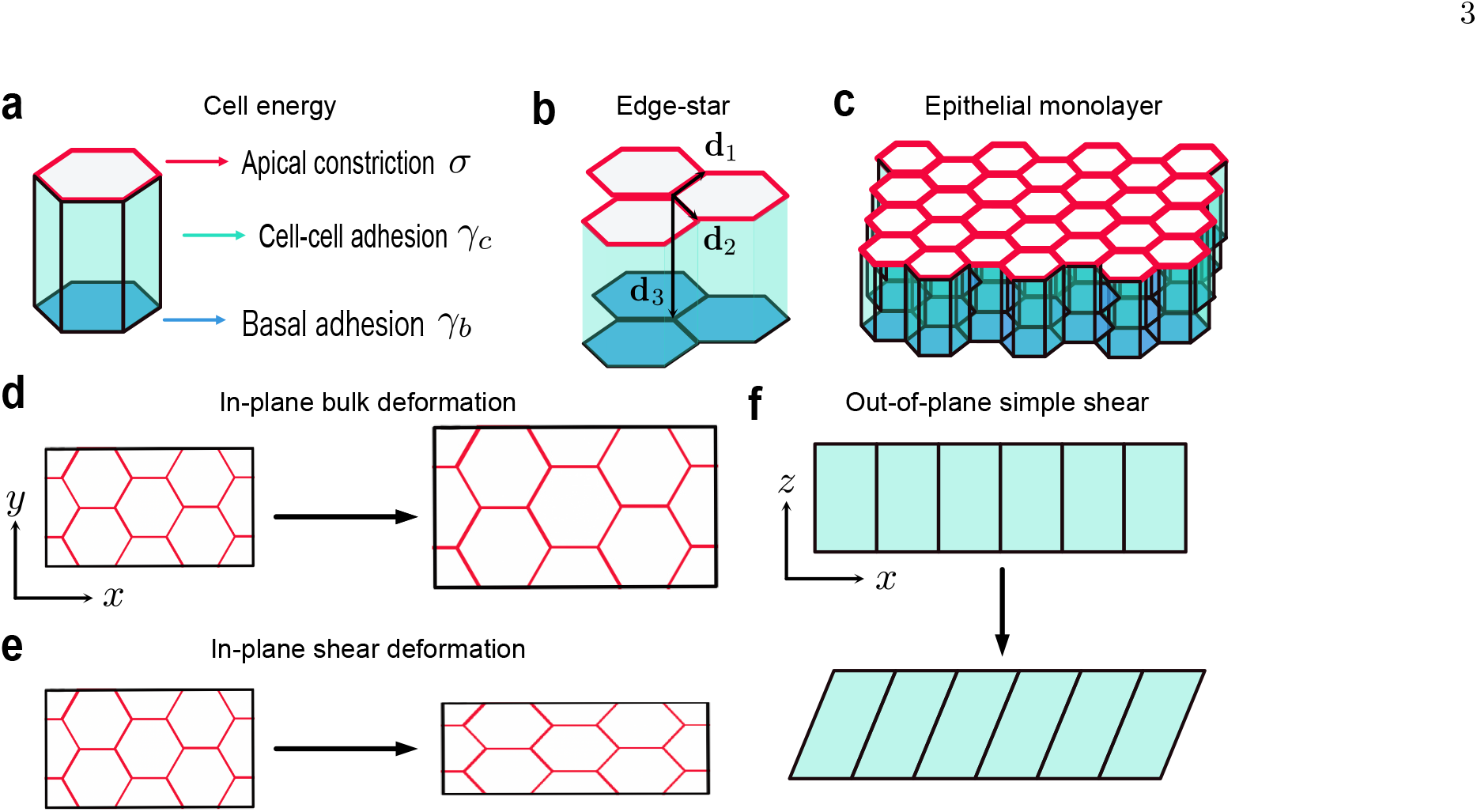
Schematic of our analysis. (a) Energy contributions of a prismatic cell. (b) Edge-star geometry at a vertex. (c) Regular hexagonal-prism monolayer. (d–f) In-plane bulk, in-plane pure-shear, and out-of-plane simple-shear deformations, respectively.

The energy of the entire tissue is defined as follows:

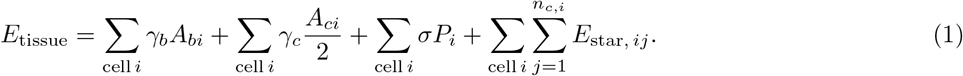

The first term is an effective basal surface-energy contribution associated with cell–substrate interactions, whereas the second is an effective lateral interfacial-energy contribution associated with cell–cell contacts. Here, *A*_*bi*_ and *A*_*ci*_ denote the basal area and the total cell–cell interfacial area of cell *i*, respectively, while *γ*_*b*_ and *γ*_*c*_ are the corresponding surface energy densities. The factor 1*/*2 in the second term avoids double counting each cell–cell interface, which is shared by two neighboring cells. The third term represents the contractile energy of the apical actomyosin belt, where *P*_*i*_ denotes the apical perimeter of cell *i* and *σ >* 0 is the corresponding line tension. With this sign convention, negative *γ*_*b*_ favors an increase in cell–substrate contact area, whereas negative *γ*_*c*_ favors an increase in lateral cell–cell interfacial area.

This model requires a confinement energy to prevent cells from collapsing. The previous confinement energy was an inverse-square edge-length energy defined for regular hexagonal cells [17]. We extend this formulation to anisotropic deformations and non-affine vertex relaxation by introducing the edge-star term defined below. This choice is motivated by the specialized junctional architecture at tricellular contacts, where end-on actin anchoring and tension-dependent adhesion reinforcement contribute to the structural stability of cell vertices [24–26]. For cell *i* with *n*_*c,i*_ corners, we characterize the geometry at each corner *j* by

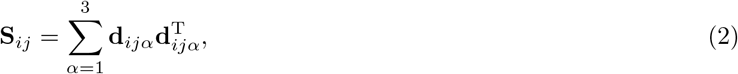

where **d**_*ijα*_ denotes one of the three incident edge vectors belonging to corner *j* of cell *i*: two edges in the apical or basal face and one apico–basal edge (Figure 1b). The corresponding edge-star energy is

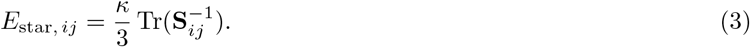

This energy penalizes strongly distorted or locally collapsed edge configurations. *κ >* 0 controls the strength of this confinement.

For a regular hexagonal prism, the energy per cell can be written as a function of the side length *l* of the apical (or basal) hexagon and the cell volume *V*_0_:

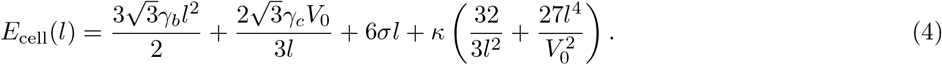

The derivation of Eq. (4) is provided in Appendix A. The equilibrium side length is obtained from *dE*_cell_*/dl* = 0.

## RESULTS

We first derive the in-plane elastic moduli and examine how three-dimensional cell configuration controls the elastic responses. We then determine how shape bistability is reflected in the elastic moduli and analyze the stability of prismatic cells against transverse shear. Finally, we formulate the mode-resolved dynamic modulus and investigate how structural disorder modifies non-affine relaxation.

### Analytical in-plane elasticity of regular prismatic cells

Here we investigate the in-plane elasticity of the model epithelium, assuming a regular hexagonal prism. The inplane elastic moduli can be computed from the change in the energy in Eq. (4) under infinitesimal deformations from an equilibrium state, following previous studies on two-dimensional cell vertex models [12, 22]. The in-plane bulk modulus *K* is obtained by considering an isotropic deformation (Figure 1d) by 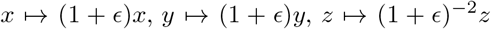 around each equilibrium state,

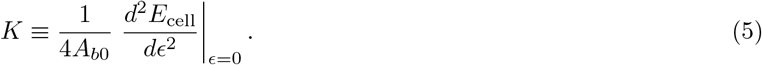

From this expression, with *A*_*b*0_ denoting area per cell in the corresponding equilibrium state, the bulk modulus *K* can be computed, resulting in

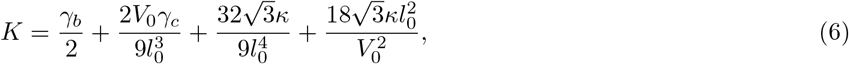

where *l*_0_ denotes the value of the cell side length *l* at the corresponding equilibrium state.

The in-plane shear modulus is obtained by considering a deformation defined by 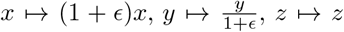 (Figure 1e),

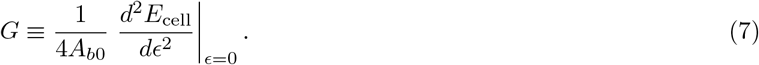

By an analogous calculation to the bulk modulus, we obtain

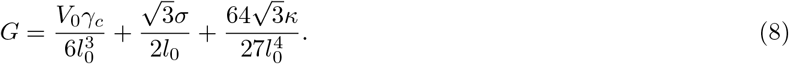

Owing to the sixfold rotational symmetry of the system in the plane, the in-plane linear elastic response is isotropic, and hence the pure-shear modulus is identical to the simple-shear modulus [27]. Using these results, the in-plane Poisson’s ratio *ν* can be calculated by [7, 27]

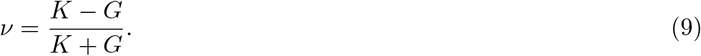

For the analyses below, we used the nondimensionalized energy function with rescaled model parameters (see Appendix B). Hereafter, we denote rescaled parameters with a tilde, e.g., 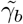for the rescaled *γ*_*b*_.

The equilibrium condition from Eq. (4) is a higher-order polynomial in the cell side length *l*, and it is thus generally difficult to obtain analytical expressions for the equilibrium configuration and the elastic moduli. However, in limiting regimes where one of basal adhesion, cell–cell adhesion, or apical constriction dominates the energy, the equilibrium cell side length, as well as the bulk and shear moduli and the Poisson’s ratio, can be determined analytically.

First, we consider the limit in which basal adhesion dominates, i.e., 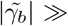1 with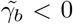. In Eq. (4), the energy contribution from basal adhesion is proportional to the square of the cell side length *l*, so in this limit, the equilibrium side length 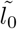diverges. Balancing the basal adhesion and edge-star terms gives 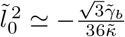Accordingly, the elastic moduli satisfy 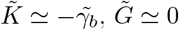and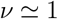. This implies that deformations that increase the epithelial sheet area are energetically costly, whereas area-preserving deformations require little energy. As a result, the cell sheet responds to in-plane stretching while approximately maintaining its thickness, corresponding to a Poisson’s ratio approaching unity. Next, we consider the limit in which cell–cell adhesion dominates, i.e.,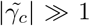 1 with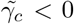. Following the same procedure, we obtain 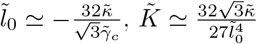 and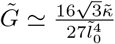 Thus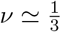

Finally, we consider the limit in which apical constriction dominates. In this regime, 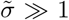, and the equilibrium cell size becomes small. Balancing the apical constriction and edge-star terms gives 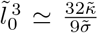The corresponding elastic moduli are 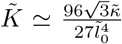 and 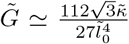 yielding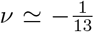. The Poisson’s ratio therefore becomes negative, indicating that under uniaxial extension from the ground state, the tissue also expands in the transverse direction. This transverse expansion can be understood from a simple geometric argument (see Appendix C). Although cells adopt the same elongated shape in both the apical-constriction- and cell–cell-adhesion-dominated limits, their Poisson’s ratios differ substantially.

### Finite-strain validation of the linear Poisson’s ratio

We numerically confirm that the in-plane Poisson’s ratio derived under infinitesimal deformations (Eq. (9)) provides a good approximation even for finite deformations. We set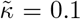.1 throughout the numerical calculations. We first fix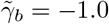.0 and determine the equilibrium side length 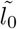by minimizing the energy of a regular hexagonal prism for each pair of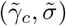. The analytical Poisson’s ratio is then calculated from the infinitesimal bulk and shear moduli as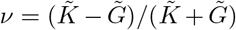. For the numerical calculation, we impose a homogeneous deformation **F** = diag(*α, β*, (*αβ*)^*−*1^), where *α* = 1 + *ϵ* and *β* are the longitudinal and transverse stretch ratios, respectively. We set *ϵ* = 0.05, minimize the deformed-cell energy with respect to *β >* 0, and denote the resulting value by *β*^*\**^. The apparent finite-strain Poisson’s ratio based on engineering strains is evaluated as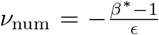. The analytical and numerical Poisson’s ratios obtained over the same 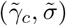 parameter space are shown in Figure 2a and 2b, respectively. The theoretical and numerical Poisson’s ratios are in good agreement over a wide range of parameters, indicating that the present theory accurately captures the elastic response of the system.

**FIG. 2.**
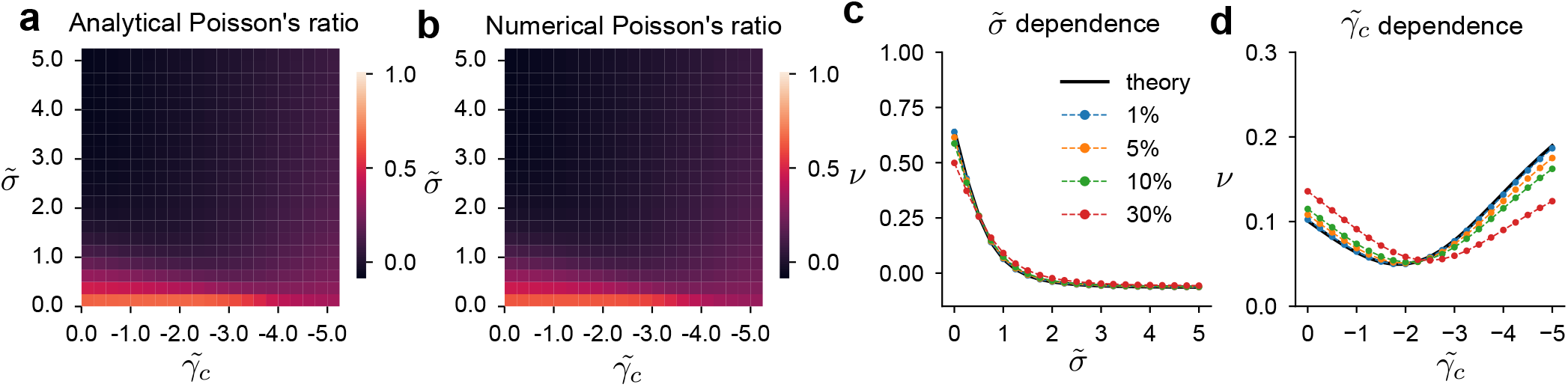
Finite-strain validation of the linear Poisson’s ratio. (a,b) Analytical and numerical Poisson’s ratios in the 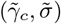 plane at 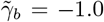panel b uses a strain of 5%. (c,d) Dependence on 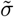at 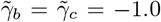and on 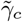at 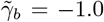and 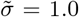respectively. Black curves show the linear theory, and symbols show numerical results at the indicated strains. Throughout, 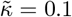

Furthermore, to assess the range of validity of the theory, we performed finite-strain numerical calculations at imposed strains of 1%, 5%, 10%, and 30%. Figure 2c shows the 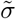-dependence of the analytical and numerical Poisson’s ratios for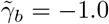.0 The theoretical prediction agrees well with the numerical results for strains up to 10% over most of the parameter range and remains close even at 30% strain, except in the low-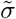 regime. Figure 2d shows the 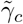-dependence for 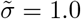and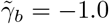. At 1% strain, the theoretical prediction agrees closely with the numerical result. Although a systematic deviation develops as the strain increases, the characteristic nonmonotonic dependence on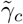, including the presence of a minimum, is retained even at 30% strain. These finite-strain deviations arise because the theory describes the linear response obtained from a quadratic expansion of the energy around the equilibrium state and therefore neglects higher-order nonlinear contributions. These results suggest that, despite its linear-response formulation, the theory captures the principal behavior of the Poisson’s ratio over the range of deformations examined here, up to 30%.

### In-plane elastic responses of prismatic cells

Having established the in-plane elastic response of the regular prismatic state, we next examine how its equilibrium structure and mechanical properties vary across the parameter space. The system can be either monostable or bistable, consistent with experimentally observed epithelial shape bistability [17, 28, 29]. Figure 3a shows the stability phase diagram at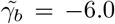. Figure 3b illustrates how the energy landscape changes from having a single minimum to having two minima as the parameter, 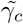is varied. To clarify their elastic responses, we analyze the monostable and bistable regimes in detail.

**FIG. 3.**
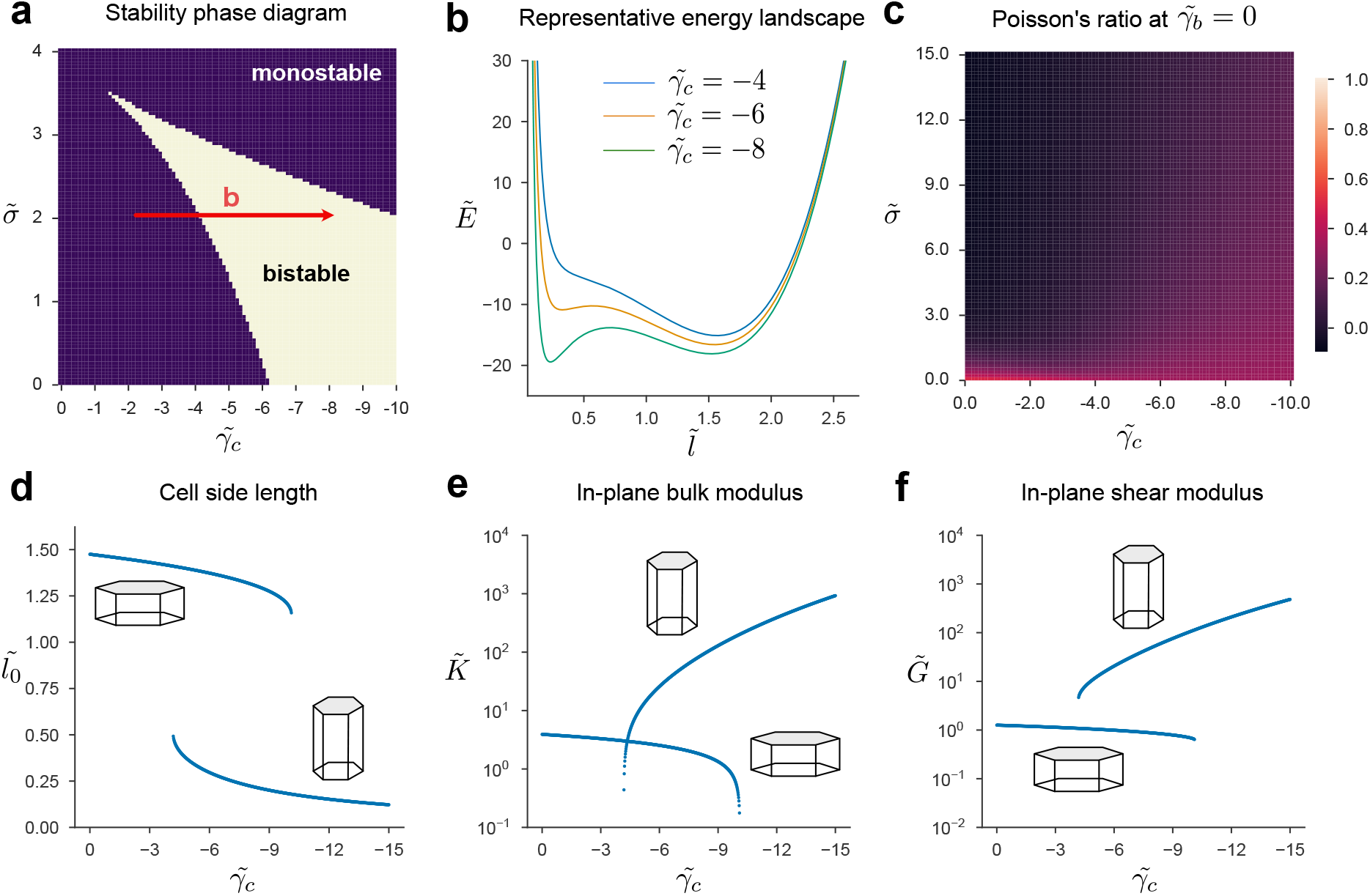
In-plane mechanics of regular prismatic cells. (a) Mono-/bistable phase diagram at 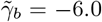.(b) Energy profiles at the indicated 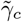along 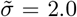.(c) Poisson’s ratio at 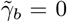.(d–f) Bifurcation diagrams of the equilibrium side length, bulk modulus, and shear modulus at 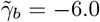and 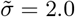,respectively. Throughout, 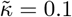

First, we consider the case 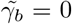and 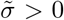in which the system is always monostable. Setting 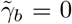provides a minimal representation of a suspended monolayer without cell-substrate adhesion [6, 30]. The in-plane Poisson’s ratio, defined in Eq. (9), approaches zero over a wide range of parameters as shown in Figure 3c. Importantly, this behavior has also been observed in actual suspended cell monolayers [6]. Moreover, in the two-dimensional model analyzed in Ref. [23], a zero Poisson’s ratio occurs only on a restricted parameter set. These results suggest that three-dimensional cell geometry together with volume conservation plays a key role in the elastic response of epithelial tissues.

When the system is bistable, the bifurcation of the equilibrium cell shape is accompanied by a bifurcation of the elastic moduli. Figures 3d–f show bifurcation diagrams of the equilibrium cell side length, in-plane bulk modulus, and in-plane shear modulus, respectively, as a function of 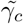at 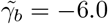and 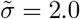In the squamous-cell branch, corresponding to the upper branches in Figure 3d and the lower branch in Figures 3e and f, basal adhesion makes a substantial contribution to the energy. As a result, the bulk modulus is several times larger than the shear modulus (Figure S1). In contrast, along the columnar-cell branch, the bulk modulus is approximately twice the shear modulus, which means the Poisson’s ratio approaches 1*/*3 (Figure S1). This behavior is consistent with the limiting behavior dominated by cell–cell adhesion.

### Affine out-of-plane shear stability of regular prismatic cells

Next, we examine the response to out-of-plane simple shear, in which the shear plane contains the direction normal to the epithelial sheet. The out-of-plane shear modulus *G*_*⊥*_ is obtained by considering an out-of-plane simple shear deformation (Figure 1f) by *x → x* + *qz, y → y, z → z* around each equilibrium state,

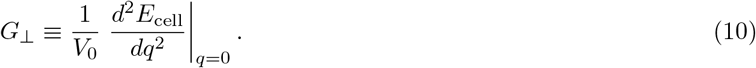

We restrict the analysis to affine deformations and examine the out-of-plane shear response of the regular hexagonal prism configuration. Under an out-of-plane simple shear, the energy per cell can be written as

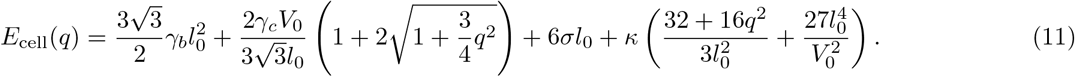

Using Eq. (10), we obtain the out-of-plane shear modulus

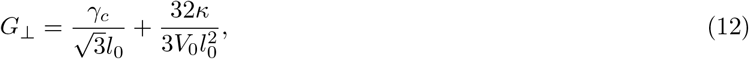

where *l*_0_ denotes the equilibrium cell side length within the regular-hexagonal-prism subspace. For *G*_*⊥*_ *>* 0, the regular hexagonal prism configuration at *q* = 0 is locally stable against out-of-plane shear, whereas for *G*_*⊥*_ *<* 0, the *q* = 0 state is locally unstable. In the latter case, the energy landscape restricted to affine deformations takes a double-well form, as illustrated by the orange curves in II and III of Figure 4b, and stable states emerge at finite shear strains *q* = *±q*_0_.

**FIG. 4.**
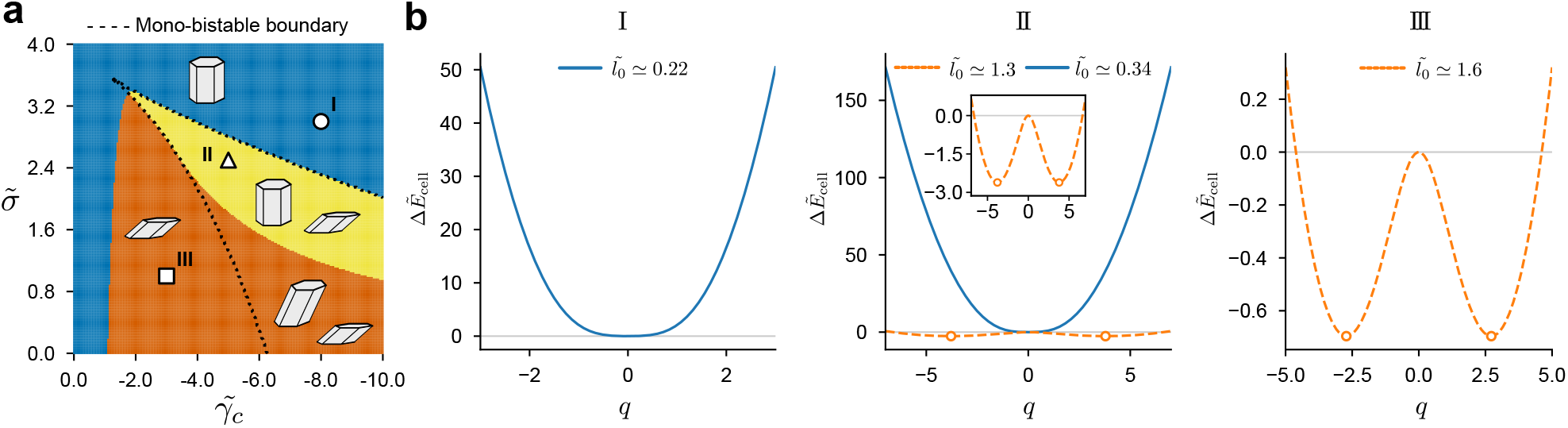
Affine out-of-plane shear stability of regular prismatic cells. (a) Stability diagram at 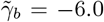and 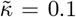 Blue, yellow, and orange indicate regions with only *q*_0_ = 0, coexistence of *q*_0_ = 0 and *q*_0_ = 0, and only *q*_0_ = 0, respectively. Dotted curves show the mono-/bistable boundary. (b) Energy landscapes at I (8.0, 3.0), II (5.0, 2.5), and III (3.0, 1.0), where the coordinates denote 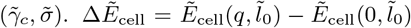 At II, the two curves correspond to the two in-plane branches; circles mark finite-shear minima.

Within the affine-deformation subspace, the equilibrium states at 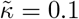1and 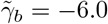0can be classified into three regimes according to their stability against out-of-plane shear (Figure 4a). In the blue region, all stable equilibrium states remain unsheared (*q* = 0) (Figure 4b, I), whereas in the yellow region, an unsheared equilibrium state coexists with stable states at finite shear (Figure 4b, II). In the orange region, only the finite-shear equilibrium states are stable (Figure 4b, III).

The out-of-plane pure shear modulus is given by 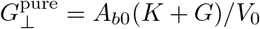and remains positive over the parameter range examined here. Together, these results establ*⊥*ish how three-dimensional cell geometry and shape bistability govern the static mechanical response of regular prismatic epithelia, from in-plane elasticity and Poisson’s ratio to stability against out-of-plane shear.

### Hessian-mode formulation of linear dynamic moduli

We then turn to the dynamic properties of the model by examining its frequency-dependent response. We denote the externally imposed affine deformation from an equilibrium state by *ϵ* and the resulting non-affine displacement by **u**, and assume overdamped dynamics of the form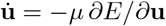. Expanding *E* to second order in **u** and *ϵ* about the equilibrium state, we define the Hessian matrix **H**, the deformation–displacement coupling vector **Ξ**, and the affine elastic coefficient *C* as

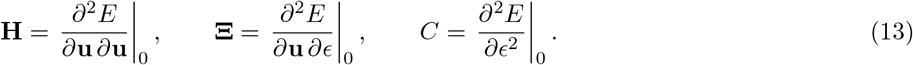

Here, the subscript 0 denotes evaluation at the equilibrium state. For an oscillatory affine deformation, *ϵ*(*t*) = *ϵ*_0_*e*^*iωt*^, the linear dynamic modulus is given by

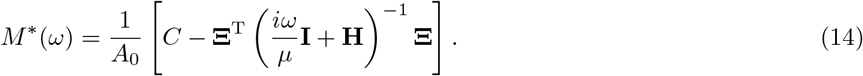

The strain parameter *ϵ* denotes the infinitesimal areal strain for bulk deformation and the engineering shear strain for simple shear; these conventions account for the absence of the factor 1*/*4 used with the axial-strain parameter in the static calculations above.

To interpret this expression, we decompose the response into the eigenmodes **e**_*j*_ of the Hessian **H**, defined by **He**_*j*_ = *h*_*j*_**e**_*j*_:

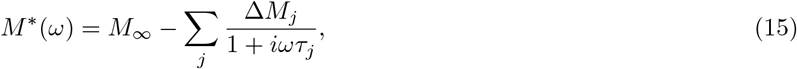

where *M*_*∞*_ = *C/A*_0_ is the high-frequency affine modulus, *A*_0_ is the equilibrium in-plane area of the periodic simulation box, and *τ*_*j*_ = 1*/*(*µh*_*j*_) is the relaxation time associated with Hessian eigenmode *j*. The relaxation strength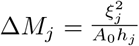 quantifies the contribution of mode *j* to the relaxation of the modulus, where 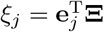 measures the coupling between the imposed affine deformation and mode *j*. The sums are taken over the nonzero eigenmodes of the Hessian.

Writing the dynamic moduli as *M*^*\**^(*ω*) = *M*^*′*^(*ω*) + *iM*^*′′*^(*ω*), the storage and loss moduli are given by

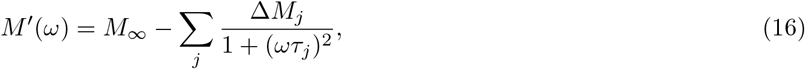

and

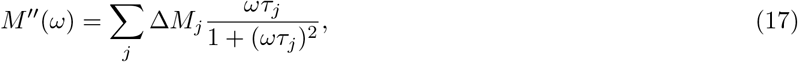

respectively. Therefore, the contribution of each eigenmode *j* to the loss modulus is maximized when *ωτ*_*j*_ = 1, or equivalently, when *ω* = *µh*_*j*_. A detailed derivation is provided in Appendix D.

### Structural disorder modifies non-affine relaxation

To examine how structural disorder modifies the response beyond the regular affine description, we applied the linear-response framework above to regular hexagonal-prism and disordered tilings under oscillatory bulk and simple shear deformations. We used 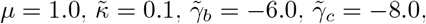 and 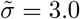 and compared configurations containing *N*_cell_ = 40 cells in the same periodic box. We retained apico-basal symmetry and a common cell height, with two independent in-plane degrees of freedom per vertex. For these calculations, individual cell volumes were constrained by a soft volume penalty rather than an exact incompressibility constraint. The generation and equilibration of the disordered tilings are described in Supplementary Material S1. For the normalized quantities shown below, we use the median nonzero Hessian eigenvalue of the regular tiling, *h*_0_, as a reference scale and define *τ*_0_ = (*µh*_0_)^*−*1^ and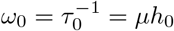.

The frequency-dependent response of the regular tiling distinguishes the affine elastic response from relaxation through internal vertex motions. At high frequencies, the storage moduli agreed with the analytical affine bulk and shear moduli derived above (Figures 5a and 5b). The bulk storage modulus remained independent of frequency, and the bulk loss modulus vanished (Figures 5a and 5d). In contrast, the shear storage modulus decreased toward low frequencies, accompanied by a finite shear loss modulus (Figures 5b and 5e). Thus, non-affine shear relaxation was present even in the regular tiling.

**FIG. 5.**
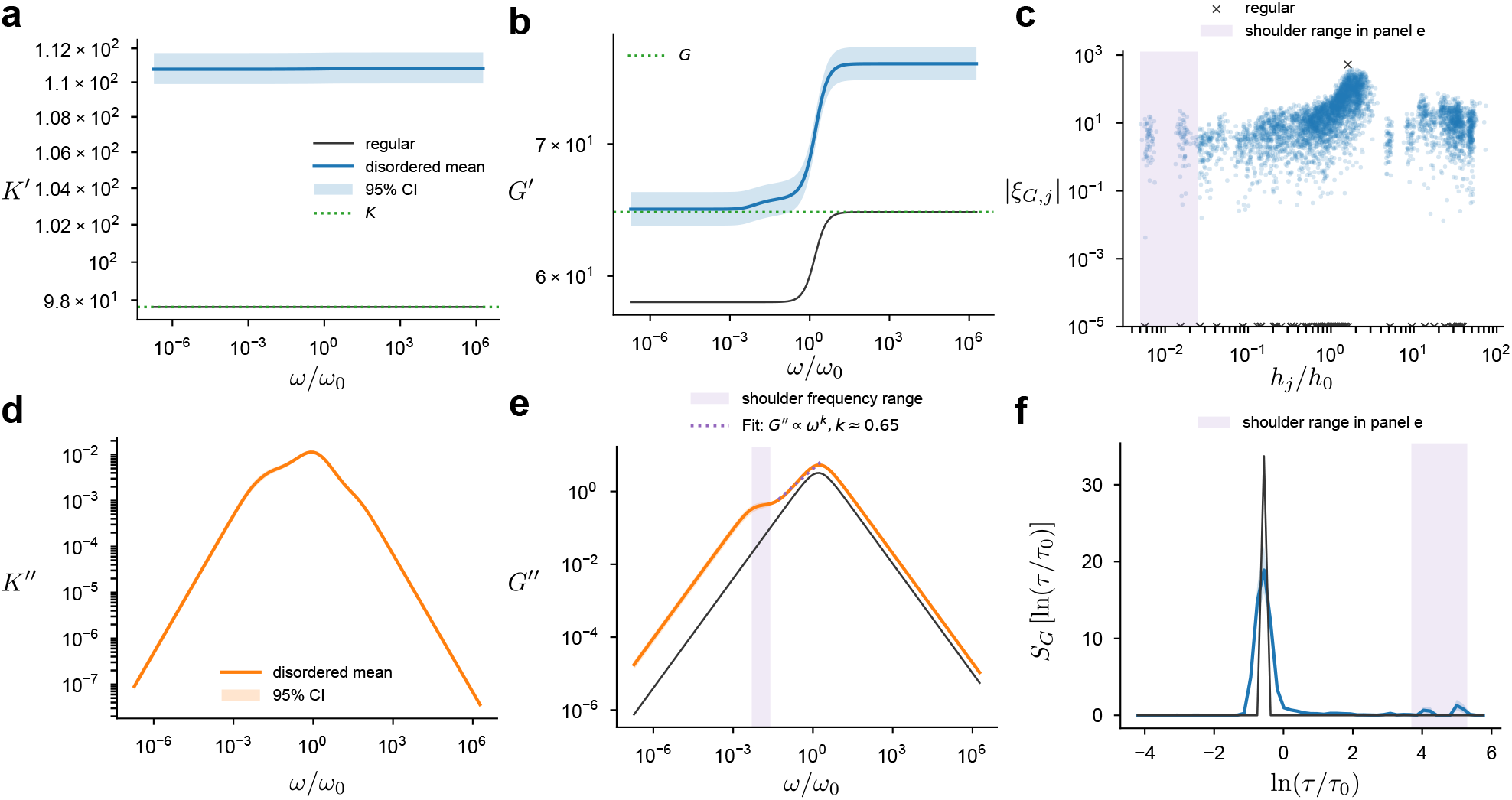
Dynamic response of regular and disordered tilings. (a,d) Bulk storage and loss moduli. (b,e) Simple shear storage and loss moduli. Colored curves show the mean over 30 disordered tilings, with shaded bands indicating pointwise 95% confidence intervals obtained by percentile bootstrap with 5,000 resamples of the tilings. Black curves show the regular tiling; green dotted lines show the analytical affine moduli. The regular bulk loss modulus vanishes. The purple dotted line in (e) shows a power-law fit, *G*^*′′*^*∝ ω*^*k*^, to the disordered ensemble mean over 0.05 *≤ω/ω*_0_ *≤*2.5. (c) Coupling of the shear deformation to Hessian modes, pooled over all realizations. (f) Shear relaxation spectrum evaluated using 50 equal-width bins in ln *τ*. The value in each bin *k* is *S*_*G,k*_ = (▵ ln *τ*)^*−*1^ _*j∈*bin *k*_ ▵*G*_*j*_, where 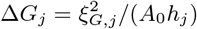 is the mode relaxation strength and ▵ ln *τ* is the bin width. Eigenvalues, relaxation times, and frequencies are normalized by *h*_0_, *τ*_0_ = (*µh*_0_)^*−*1^, and *ω*_0_ = *µh*_0_, respectively, where *h*0 is the median nonzero Hessian eigenvalue of the regular tiling. S2.

Structural disorder modifies both the elastic and dissipative components of the tissue response. For the parameter set examined, the disorder-averaged bulk and shear storage moduli exceeded those of the regular tiling throughout the sampled frequency range (Figures 5a and 5b). Both decreased from their respective high-frequency limits as non-affine relaxation became effective at lower frequencies, although the frequency dependence of the bulk storage modulus was weak. Unlike the regular tiling, the disordered tilings exhibited a finite bulk loss modulus, indicating that bulk deformation coupled to non-affine vertex motions (Figure 5d). The mean shear loss modulus was also larger than that of the regular tiling and exhibited a shoulder on the low-frequency side of the main loss peak (Figure 5e, shaded region).

To examine the modal origin of this shoulder, we analyzed the coupling of shear deformation to Hessian modes |*ξ*_*G,j*_| and the associated relaxation spectrum,

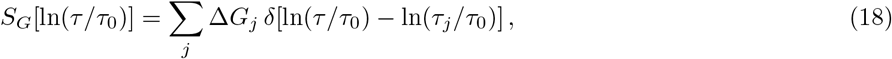

where ▵*G*_*j*_ is the relaxation strength of mode *j*. For the regular tiling, appreciable shear coupling was concentrated at a single Hessian eigenvalue, whereas the disordered tilings coupled to modes spanning a broader range of eigenvalues (Figure 5c). Correspondingly, the relaxation spectrum of the regular tiling exhibited a sharp peak, while that of the disordered tilings was broader and contained additional spectral weight at longer relaxation times, including near ln(*τ/τ*_0_)*≈* 5 (Figure 5f). Since each mode contributes most strongly to the loss modulus at *ωτ*_*j*_ = 1, these slow, shear-coupled modes contribute to the low-frequency shoulder. The shaded ranges in Figures 5c and 5f identify modes whose loss maxima fall within the shoulder frequency range highlighted in Figure 5e. The contributions of these modes give rise to a regime in which *G*^*′′*^ increases more slowly with frequency and is well described by the sublinear scaling *G*^*′′*^*∝ ω*^0.65^ (the purple dotted line in Figure 5). Thus, disorder broadens the range of relaxation times contributing to the shear response, with the long-time spectral weight providing a modal interpretation of the shoulder. Both the shoulder and the long-time relaxation components persisted across the cell numbers and volume penalties examined in Figures S3 and S4.

The corresponding complex Poisson’s ratio, *ν*^*\**^(*ω*) = [*K*^*\**^(*ω*) *− G*^*\**^(*ω*)]*/*[*K*^*\**^(*ω*) + *G*^*\**^(*ω*)], remained small over the frequency range examined (Figure S5). Its real part varied only moderately with frequency and remained comparable between the regular and disordered tilings, indicating that the balance between bulk and shear elasticity is preserved despite disorder-induced non-affine relaxation.

## DISCUSSION

In this study, we extend the original geometric confinement to a local regularization based on the three edges meeting at each cell corner. By penalizing configurations in which the incident edge vectors lose linear independence, the regularization accommodates anisotropic deformations and non-affine vertex motions while retaining analytical tractability for regular prismatic cells. In addition, unlike conventional two-dimensional vertex models based on target area and perimeter terms [11, 12, 22, 23], the present model explicitly incorporates basal, cell–cell, and apical mechanical contributions, making its parameters more directly interpretable in biological terms and facilitating the incorporation of additional cellular mechanisms, such as mechanosensitive E-cadherin turnover [31].

Our analysis shows that a near-zero in-plane Poisson’s ratio emerges over a broad parameter range in regular prismatic tissues. This response is not restricted to the idealized suspended-monolayer limit, *γ*_*b*_ = 0 (Figure 3), but also occurs for nonzero *γ*_*b*_ when apical constriction is sufficiently strong (Figure 2). Moreover, the complex Poisson’s ratio is only weakly altered by structural disorder over the timescales examined (Figure S5), indicating that a near-zero Poisson’s ratio is robust to structural disorder and persists across the timescales examined. These results are consistent with experimental measurements showing a small in-plane Poisson’s ratio under slow deformation [6, 7]. In particular, suspended monolayers detached from the substrate allow epithelial elasticity to be measured directly and exhibit a small in-plane Poisson’s ratio (see Fig. 3 of Ref. [6]), whereas conventional two-dimensional cell vertex models reproduce near-zero values only for restricted parameter choices [23]. The relation *ν* = (*K − G*)*/*(*K* + *G*) shows that this response reflects nearly matched in-plane bulk and shear moduli, rather than a negligible resistance to area change. Geometrically, longitudinal extension is accommodated predominantly by a reduction in cell thickness rather than by in-plane transverse contraction. Three-dimensional cell geometry together with volume conservation permits this deformation mode, while the balance among the different energy contributions determines the relative changes in width and thickness. Such a response may help epithelial sheets accommodate extension while limiting lateral contraction, a property potentially relevant to their function as continuous tissue coverings.

Allowing out-of-plane shear introduces an additional condition for the stability of these states. A regular prism that is stable against the examined in-plane deformations can nevertheless have a negative out-of-plane simple-shear modulus. Under this deformation, the increase in lateral interfacial area can lower the adhesive energy, competing with the stabilizing regularization. Unlike the in-plane strains studied in strictly planar vertex models [11, 12, 22, 23], this mode involves relative displacement of the apical and basal surfaces and requires a description that resolves deformation across the tissue thickness. Experiments on epithelial curling demonstrate that out-of-plane deformation and in-plane tension can be mechanically coupled [32]. It would therefore be interesting to extend the present analysis to include tissue bending and examine whether the identified shear instability contributes to epithelial curling.

Extending the model through a local regularization of cell-corner geometry provides a route for linking heterogeneous tissue structure to both viscoelastic responses and experimentally inferred mechanical parameters. In our simulations, structural disorder broadens the spectrum of non-affine relaxation modes and introduces additional slow contributions to the frequency-dependent response (Figure 5). These qualitative features resemble those reported in two-dimensional vertex models [15, 33–35], despite the three-dimensional origin of the present energy and volume constraints. For quantitative comparison with experiments, however, the model parameters must also be inferred from data. Recent advances in differentiable simulation have begun to enable parameter inference in cell-based models directly from imaging data [36, 37]. Applying such approaches to three-dimensional cell vertex models could make it possible to infer spatially heterogeneous mechanical parameters and test how this heterogeneity shapes tissue-scale elastic and viscoelastic responses.

Further theoretical and experimental work could assess the robustness of the present predictions. The form of the local regularization is not uniquely prescribed, and the sensitivity of the elastic moduli and stability boundaries to alternative constitutive choices remains to be examined. Incorporating cortical turnover and mechanochemical feed-back would also allow adhesion and contractility to evolve rather than being represented by fixed coefficients [38, 39]. Experimentally, the model predicts that the monolayer enters a near-zero Poisson’s-ratio regime as apical contractility becomes dominant. This qualitative prediction could be tested by combining optogenetic control of apical constriction [40] with small-strain tensile measurements. Measuring incremental longitudinal and transverse strains together with thickness changes would reveal how extension is partitioned between transverse deformation and thinning.

## CONCLUSION

We extended a three-dimensional cell vertex model of epithelial monolayers to accommodate non-affine vertex motions and structurally disordered configurations through a local regularization of cell-corner geometry. Using this framework, we analytically characterized the in-plane elastic response of regular prismatic cells and showed that a near-zero Poisson’s ratio arises over a broad parameter range, remains robust to structural disorder, and persists across the timescales examined. Notably, states stable against the in-plane deformations examined can nevertheless be unstable to out-of-plane simple shear. Structural disorder further broadens the spectrum of non-affine relaxation modes and enhances the contribution of slow modes to the frequency-dependent response. Together, these results provide a unified framework for relating three-dimensional cell geometry, elastic stability, and internal relaxation in epithelial tissues.

## APPENDIX A ENERGY OF THE REGULAR HEXAGONAL TILING AND AFFINE DEFORMATIONS

For a regular hexagonal prism with side length *l* and height *h*, the basal area, total lateral area, and apical perimeter are

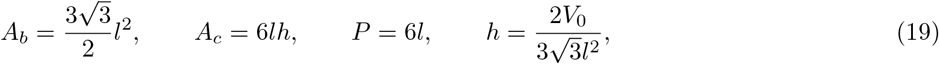

where the last relation follows from volume conservation, *A*_*b*_*h* = *V*_0_.

At each corner of a cell, two horizontal edges of length *l* and one apico–basal edge of length *h* meet. The two horizontal edges form an angle of 120^*°*^. In a local frame whose in-plane axes lie along their angle bisectors, the corner tensor is

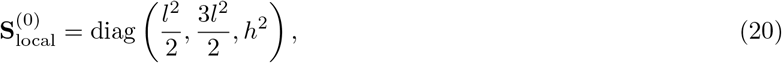

and hence

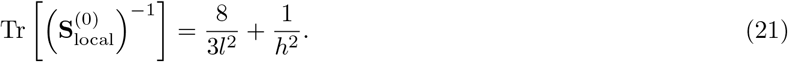

All twelve corners of a hexagonal prism contribute, without an additional vertex-sharing factor. The edge-star energy per cell is therefore

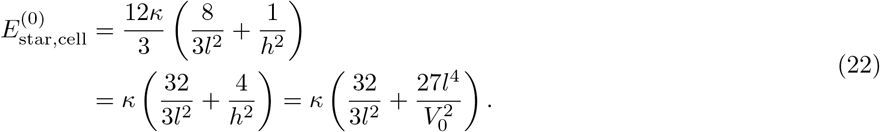

Combining the four energy contributions gives

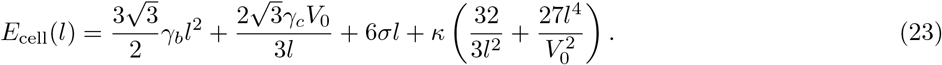

Under a homogeneous affine deformation **F**, each incident edge vector transforms as **d**_*ijα*_ *→* **Fd**_*ijα*_. Thus,

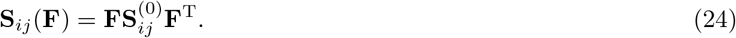

Individual corner tensors are anisotropic in the plane. Expressing their inverses in a common frame and summing over the twelve corners, sixfold symmetry gives

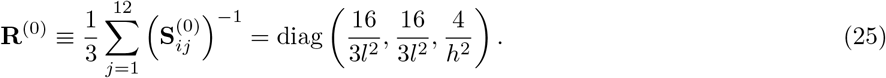

The affine edge-star energy is therefore

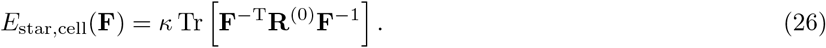

Here, **R**^(0)^ is a sum of inverse tensors, not the inverse of their sum. Substitution of the bulk, in-plane pure-shear, and out-of-plane simple-shear deformation gradients used in the main text gives the corresponding affine edge-star energies.

## APPENDIX B NONDIMENSIONALIZATION

We define the characteristic length using the prescribed cell volume,

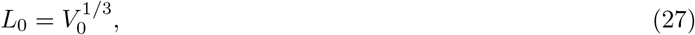

and denote the reference energy used in the numerical calculations by *E*_ref_. Dimensionless geometric quantities are defined by

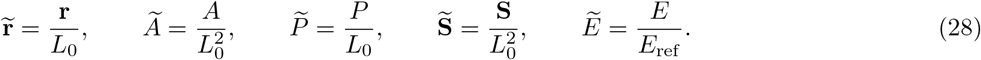

The dimensionless model parameters are

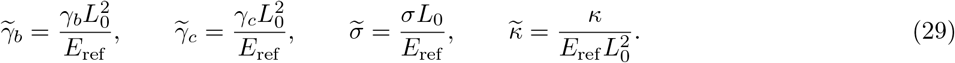

Defining 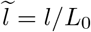the dimensionless energy of a regular hexagonal prism is

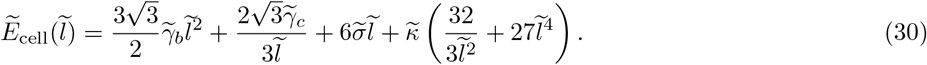

The in-plane and out-of-plane elastic moduli are rescaled as

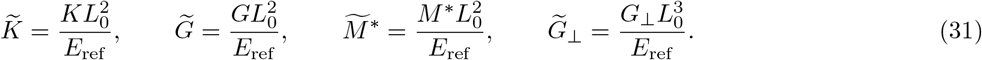

The Poisson’s ratio is dimensionless and is unchanged by this rescaling. Unless otherwise stated, tildes denote the dimensionless quantities defined above.

## APPENDIX C MECHANISM OF NEGATIVE POISSON’S RATIO IN THE APICAL-CONSTRICTION-DOMINANT LIMIT

In the limit of strong apical constriction, the leading contributions to the energy of a regular prism are

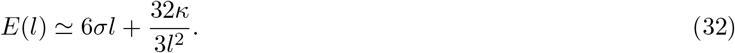

The equilibrium condition then gives

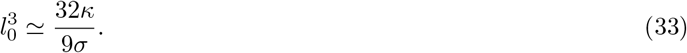

We consider a small homogeneous in-plane deformation

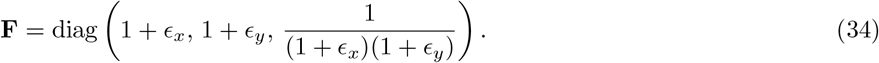

The apical perimeter of the deformed regular hexagon is

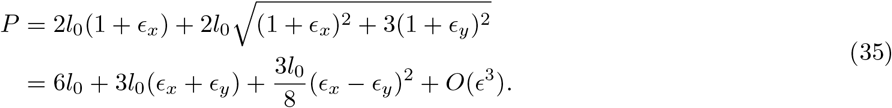

The leading in-plane contribution to the edge-star energy is

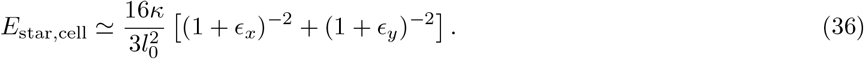

The basal, lateral, and vertical edge-star contributions are subleading in the limit *l*_0_→ 0. Expanding the energy to second order gives

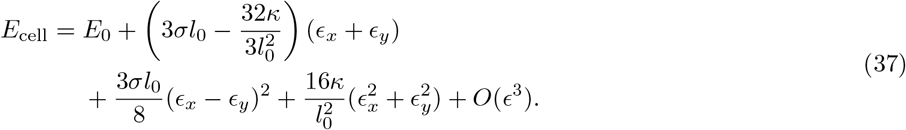

Using the equilibrium relation, the linear term vanishes and the quadratic energy becomes

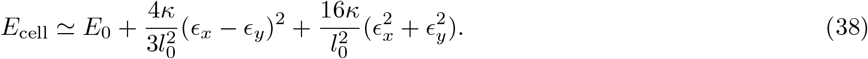

Minimizing this expression with respect to *ϵ*_*y*_ at fixed *ϵ*_*x*_ gives

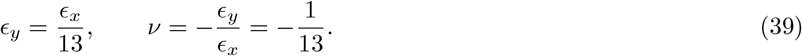

Thus, the transverse direction expands under longitudinal extension, producing a negative in-plane Poisson’s ratio.

## APPENDIX D DETAILED DERIVATION OF THE LINEAR DYNAMIC MODULUS

Expanding the energy around an equilibrium state by **u** and *ϵ* gives

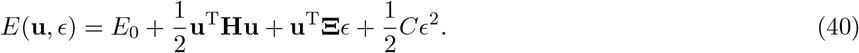

For overdamped dynamics,

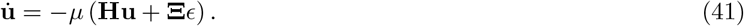

Under an oscillatory strain 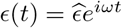the steady-state displacement satisfies

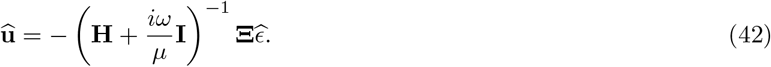

Denoting the generalized stress by Σ, its oscillatory component is

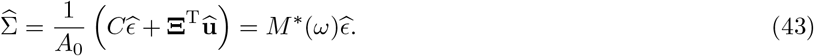

The complex modulus is therefore

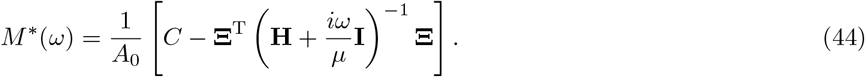

Let **He**_*j*_ = *h*_*j*_**e**_*j*_ and define

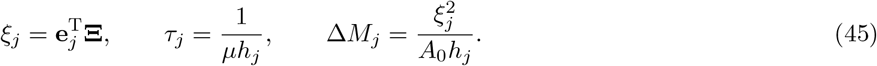

The sums below are taken over the nonzero Hessian modes. The complex modulus can then be written as

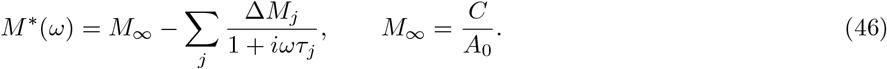

Consequently, the storage and loss moduli are

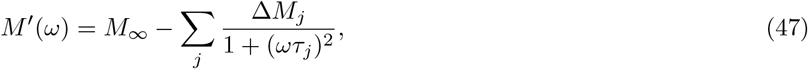

and

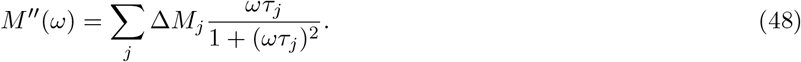

The quasistatic modulus is *M*_0_ = *M*_*∞*_ *−* _*j*_ ▵*M*_*j*_, and the contribution of mode *j* to the loss modulus is maximal at

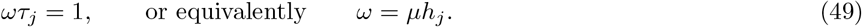

## ACKNOWLEDGMENTS

This research was supported by Japan Science and Technology Agency PRESTO (JPMJPR25K3 to Y.K.) and JSPS KAKENHI (JP25K09578 and JP26H00458 to Y.K.).

## AUTHOR CONTRIBUTIONS

Y.K. designed the research. K.T. performed theoretical analyses and numerical computations. K.T. and Y.K. wrote the manuscript.

## DATA AVAILABILITY

No experimental data were generated or analyzed in this study.

## CODE AVAILABILITY

The code used for the numerical simulations is publicly available at https://github.com/KodaiTerada/mechanics-3D-CVM.

## SUPPLEMENTARY MATERIAL

### S1. Generation of Disordered Tilings and Relaxation

We generated 30 independent disordered tilings containing *N*_cell_ = 40 cells (random seeds 100–129). Each tiling occupied the same rectangular periodic box as the regular reference, with 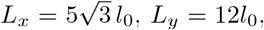and *A*_0_ = *L*_*x*_*L*_*y*_, where *l*_0_ minimizes the regular-prism cell energy. Seed points were drawn uniformly and accepted sequentially only if their minimum-image distance from every previously accepted point was at least

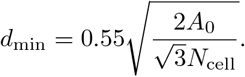

We constructed the periodic Voronoi tessellation and applied ten Lloyd iterations, moving each seed to its cell’s area centroid and recomputing the tessellation at each iteration. The polygons were extruded into prisms with identical apical and basal faces and a common height *h* = *V*_tot_*/A*_0_, where *V*_tot_ = *N*_cell_*V*_0_. The box and total volume were fixed during relaxation, and only the in-plane vertex coordinates were varied.

We first minimized the tissue energy without a cell-wise volume penalty, allowing individual cell volumes *V*_*i*_ = *A*_*bi*_*h* to change. An initial Adam stage used up to 8000 iterations with learning rate 5*×* 10^*−*5^ and allowed T1 neighbor exchanges for edges of dimensionless length at most 5*×*10^*−*3^. For each eligible edge, the exchanged and unchanged-connectivity candidates were reopened to 1.1 times this cutoff; a valid exchanged configuration was accepted if its energy did not exceed that of the unchanged-connectivity candidate. We then fixed the topology and continued relaxation using Adam followed by BFGS. When necessary, the Adam learning rate was reduced tenfold on successive retries, and damped Hessian-based steps were used to reduce small residual forces. Updates producing self-intersecting cells were rejected.

After this zero-penalty relaxation converged, we set each cell’s relaxed volume as its target *Vi\** and added a soft volume penalty,

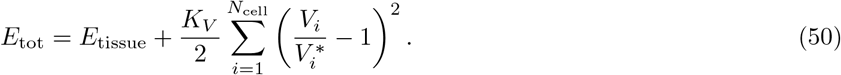

The targets and topology were held fixed during any further relaxation and the subsequent linear-response calculations.

Using the dimensionless units defined in Appendix B, we set 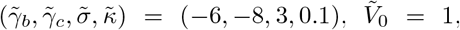 and 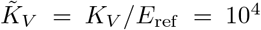. The zero-penalty and penalized relaxations were each considered converged only when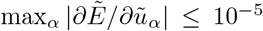, where 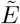 is the active energy and 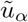 denotes an in-plane vertex coordinate. Gradients and Hessians were evaluated by automatic differentiation in double precision. Before computing linear response, we removed the two rigid-translation modes and required that the remaining Hessian have no eigenvalue below *−*10^*−*5^. Bulk and shear responses were evaluated on the same relaxed realizations.

**FIG. S1:**
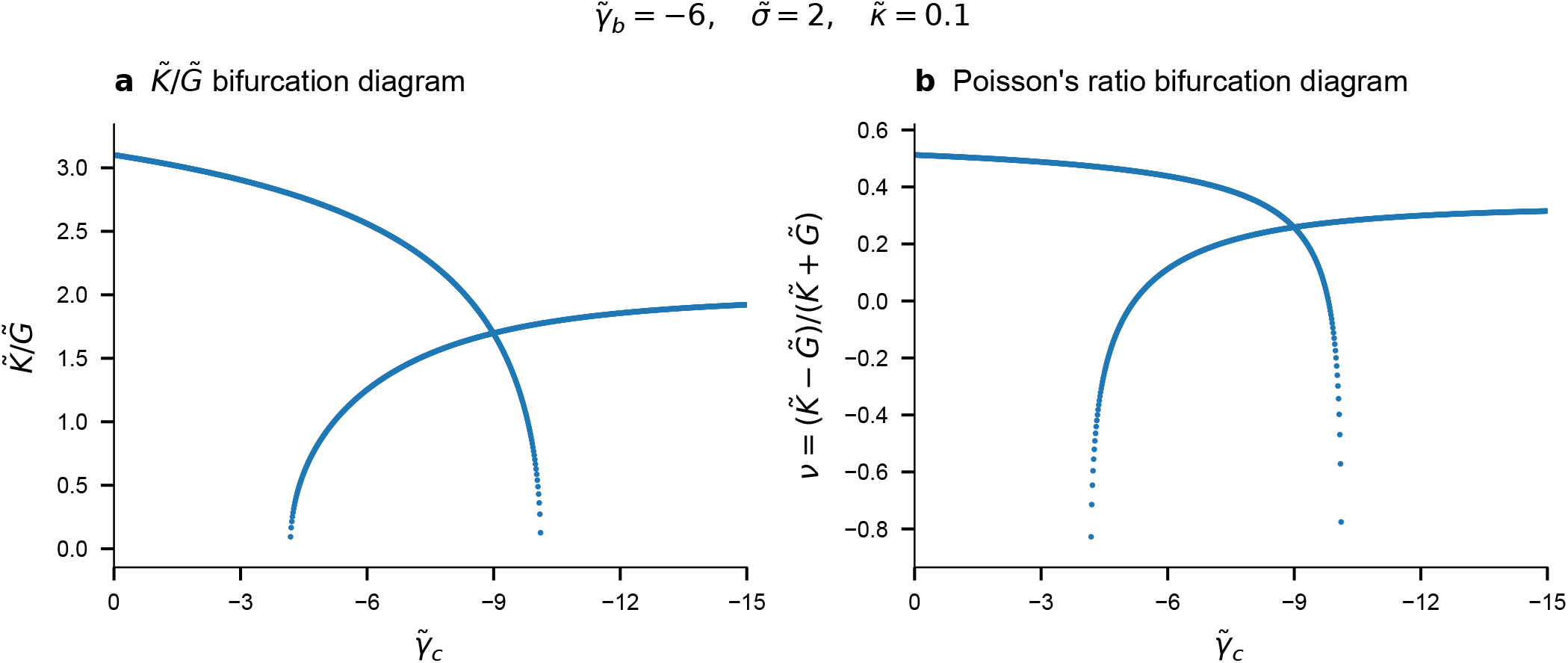
Bifurcation Diagram of 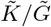 and Poisson’s Ratio. Bifurcation diagrams of the elastic-modulus ratio and Poisson’s ratio. (a) Ratio of the in-plane bulk and shear moduli, 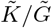and (b) Poisson’s ratio, 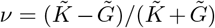as functions of 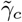at 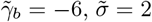and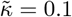. Each point corresponds to a local minimum of the regular-prism energy with respect to the cell side length, using the same branches as Figures 3d–f. Both local-minimum branches are shown in the bistable regime.

**FIG. S2.**
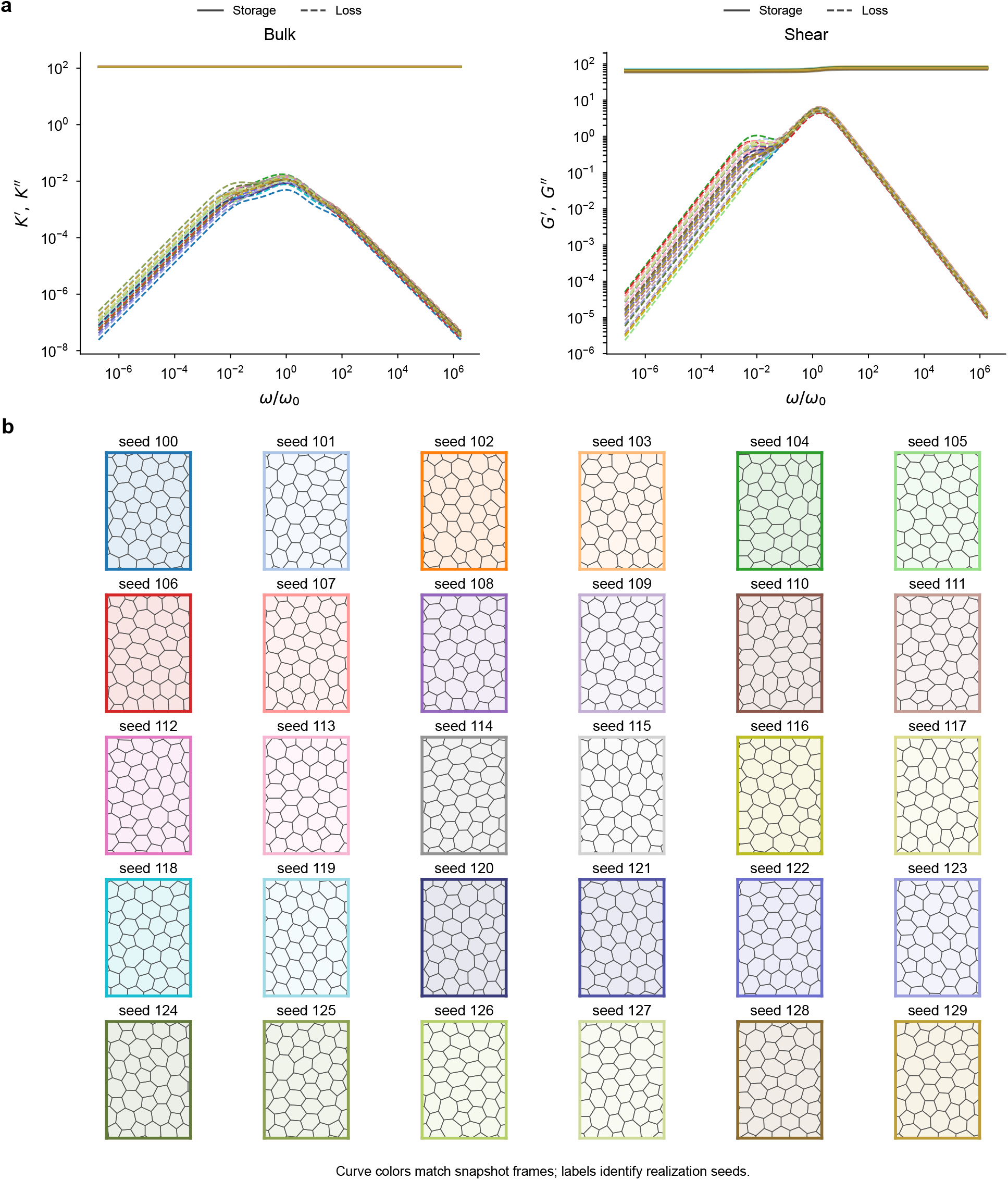
Individual realizations corresponding to Figure 5. Individual disordered realizations underlying Figure 5. (a) Bulk (left) and simple-shear (right) storage and loss moduli for the 30 disordered tilings used in Figure 5. Solid and dashed curves indicate storage and loss moduli, respectively. (b) Top views of the corresponding relaxed periodic tilings. Curve colors in panel a match the snapshot frames in panel b; labels identify the random seeds (100–129). Each tiling contains *N*_cell_ = 40 cells, with 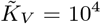 and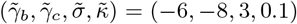. Frequencies are normalized by the regular-reference scale *ω*_0_ defined in Figure 5.

**FIG. S3.**
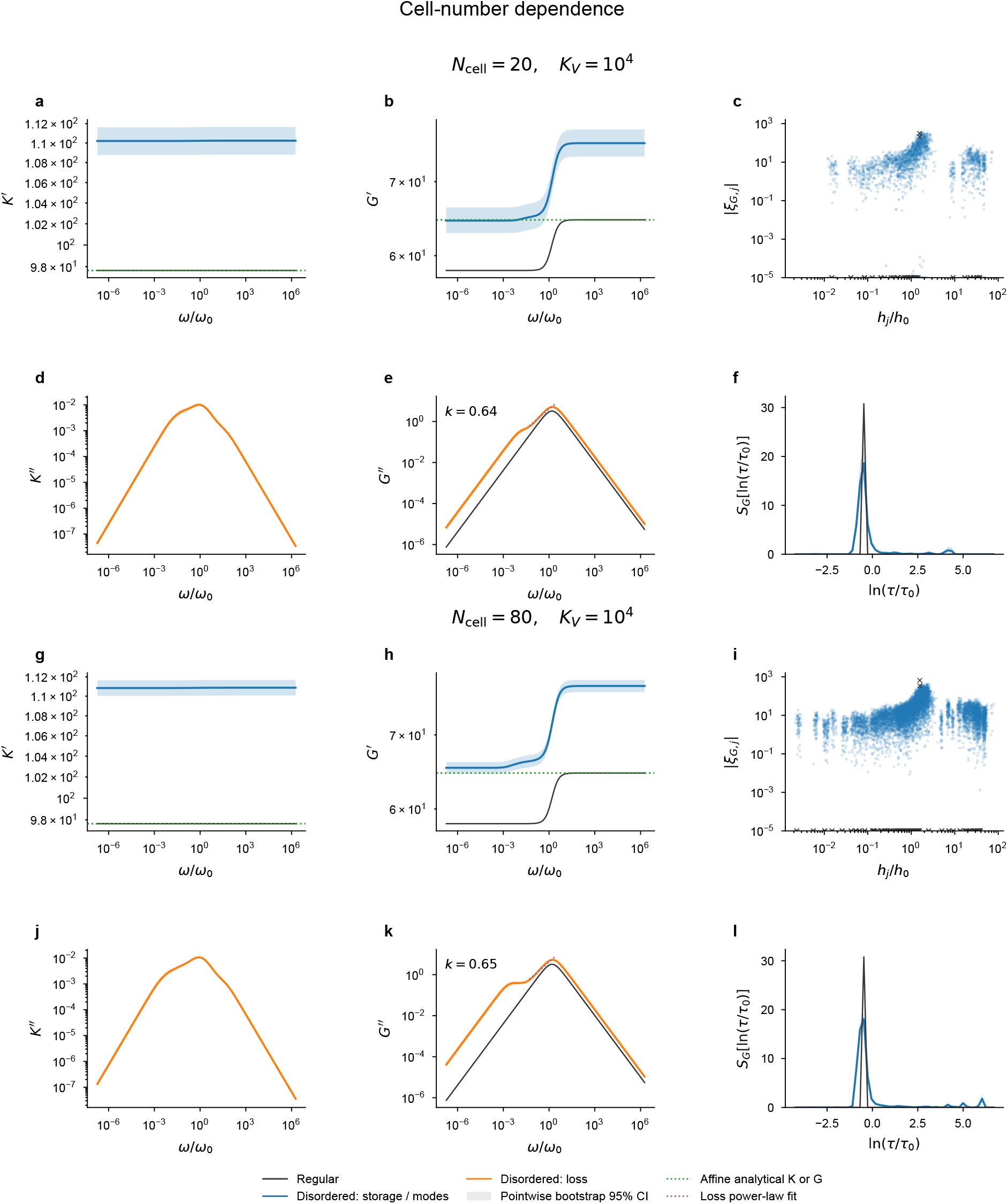
Cell-number dependence of the dynamic response. Cell-number dependence of the dynamic response. Results for *N*_cell_ = 20 (a–f) and 80 (g–l) at 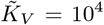Panels show bulk storage moduli (a,g), simple-shear storage moduli (b,h), shear–mode coupling magnitudes (c,i), bulk loss moduli (d,j), simple-shear loss moduli (e,k), and shear relaxation spectra (f,l). Curves and confidence intervals are defined as in Figure 5. Purple dotted lines show fits of the mean shear loss to *G*^*′′*^*∝ ω*^*k*^ over 0.05 *≤ω/ω*_0_ *≤*2.5, yielding *k* = 0.64 and 0.65, respectively. Reference scales are determined separately from the regular tiling for each condition. Throughout, 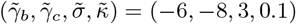 and *µ* = 1.

**FIG. S4.**
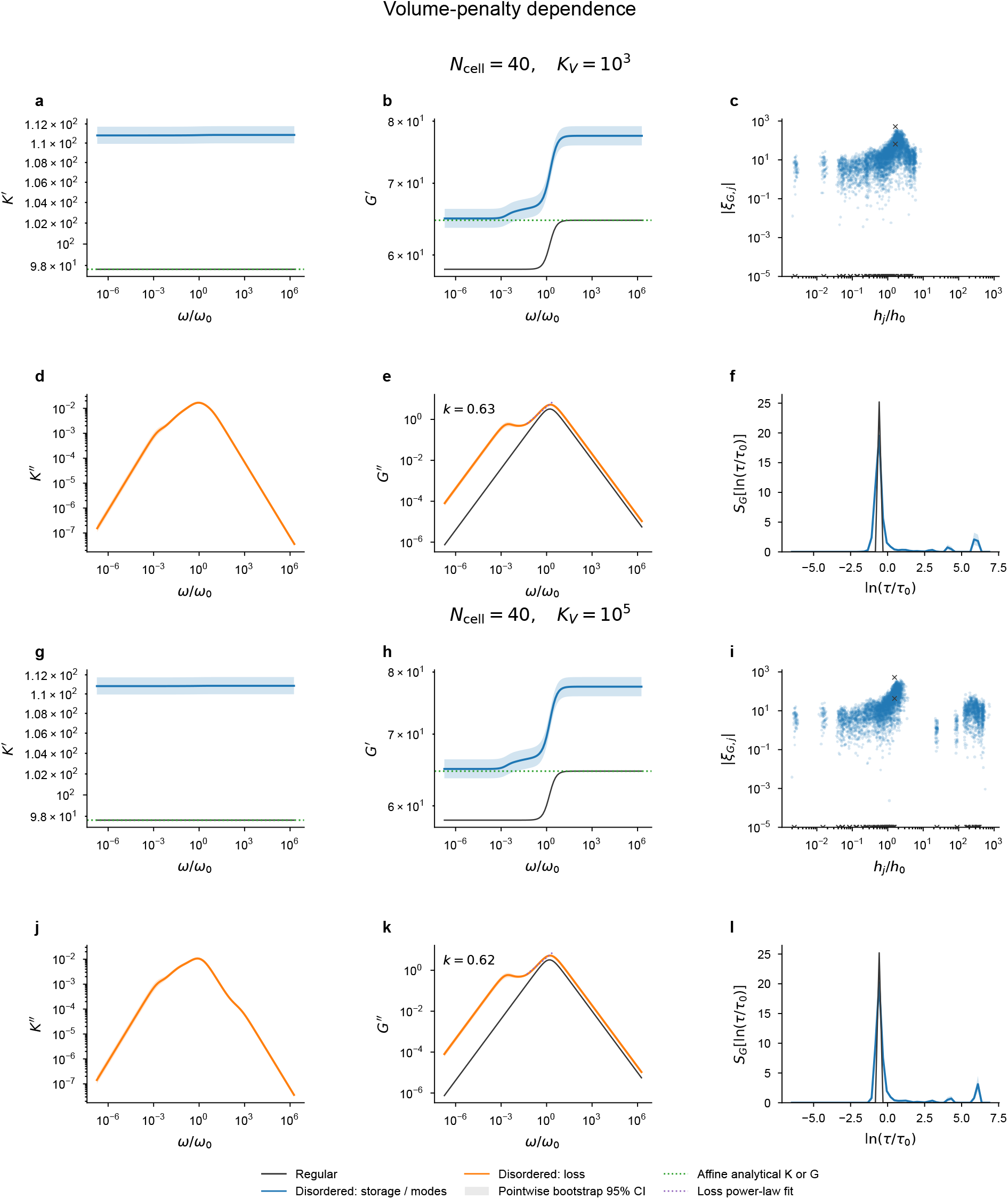
Dependence on the volume-penalty stiffness. Dependence of the dynamic response on the volume-penalty stiffness. Results for 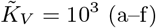and 10^5^ (g–l) with *N*_cell_ = 40; panels show bulk storage moduli (a,g), simple-shear storage moduli (b,h), shear–mode coupling magnitudes (c,i), bulk loss moduli (d,j), simple-shear loss moduli (e,k), and shear relaxation spectra (f,l). Plotting and analysis conventions follow Figure S3. The shear-loss fits give *k* = 0.63 and 0.62, respectively, and the low-frequency shoulder and long-time spectral weight persist at both penalty strengths. Throughout, 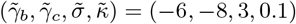and *µ* = 1.

**FIG. S5.**
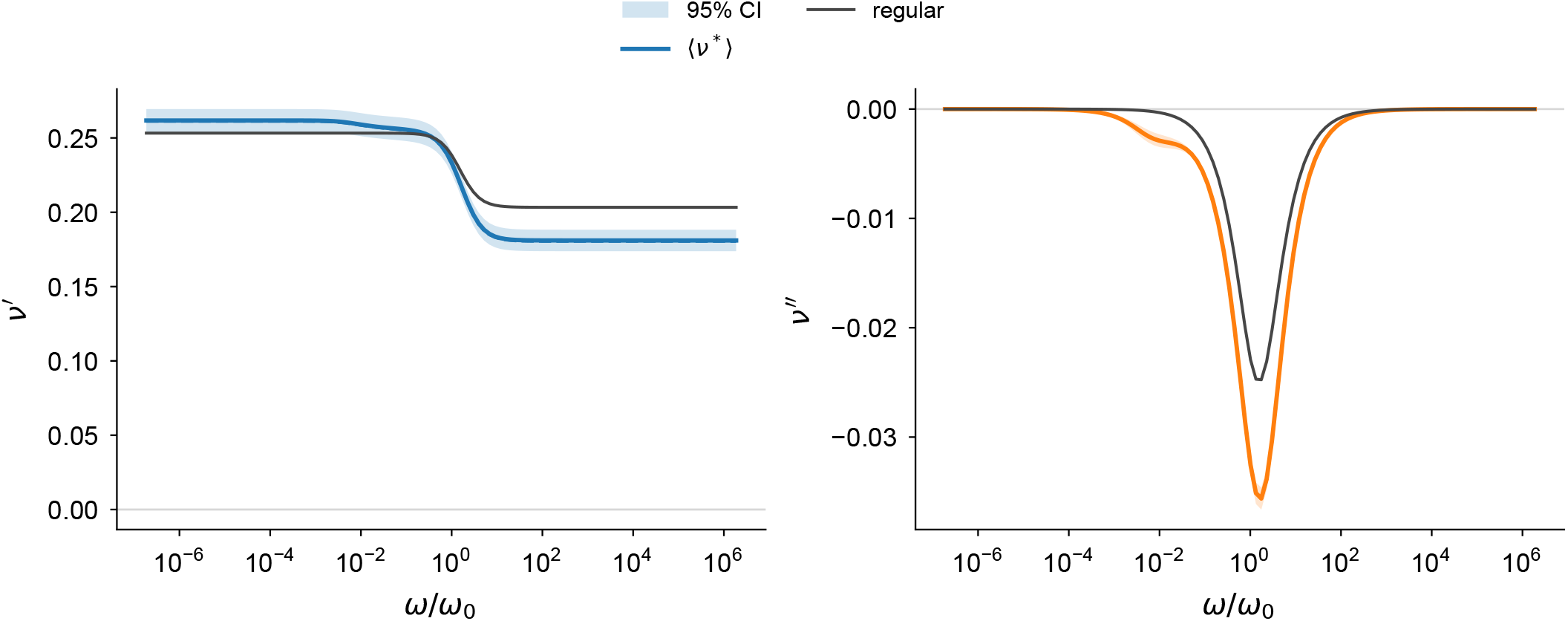
Frequency-dependent complex Poisson’s ratio. The real and imaginary parts of the complex Poisson’s ratio, *ν*^*\**^(*ω*) = [*K*^*\**^(*ω*) *G*^*\**^(*ω*)]*/*[*K*^*\**^(*ω*) + *G*^*\**^(*ω*)], are shown as functions of *ω/ω*_0_. Colored lines and shaded regions indicate the mean and 95% confidence interval over the disordered realizations, respectively, while black lines show the regular hexagonal-prism configuration. All parameters and simulation conditions are the same as those used in Figure 5. Structural disorder has little effect on the low-frequency value of *ν*^*′*^, while it lowers the high-frequency plateau and produces a more negative *ν*^*′′*^ around the characteristic relaxation frequencies.

